# Distinct attentional profile and functional connectivity of neurons with visual feature coding in the primate brain

**DOI:** 10.1101/2024.06.24.600401

**Authors:** Jie Zhang, Runnan Cao, Xiaocang Zhu, Huihui Zhou, Shuo Wang

**Affiliations:** Department of Radiology, Washington University in St. Louis, St. Louis, MO 63110, USA; Peng Cheng Laboratory, Shenzhen 518000, China; Brain Cognition and Brain Disease Institute (BCBDI), Shenzhen-Hong Kong Institute of Brain Science-Shenzhen Fundamental Research Institutions, Guangdong Provincial Key Laboratory of Brain Connectome and Behavior, CAS Key Laboratory of Brain Connectome and Manipulation, Shenzhen Institute of Advanced Technology, Chinese Academy of Sciences, Shenzhen 518055, China

**Author notes:** **Corresponding authors**: Jie Zhang,Huihui Zhou Shuo Wang. co-senior authorship (Huihui Zhou, Shuo Wang).

**Keywords:** Attention, Visual search, Object coding, Representational geometry, Spike-LFP coherence, Granger causality, V4, TE, TEO, LPFC, OFC, Macaques

## Abstract

Visual attention and object recognition are two critical cognitive functions that significantly influence our perception of the world. While these neural processes converge on the temporal cortex, the exact nature of their interactions remains largely unclear. Here, we systematically investigated the interplay between visual attention and object feature coding by training macaques to perform a free-gaze visual search task using natural face and object stimuli. With a large number of units recorded from multiple brain areas, we discovered that units exhibiting visual feature coding displayed a distinct attentional response profile and functional connectivity compared to units not exhibiting feature coding. Attention directed towards search targets enhanced the pattern separation of stimuli across brain areas, and this enhancement was more pronounced for units encoding visual features. Our findings suggest two stages of neural processing, with the early stage primarily focused on processing visual features and the late stage dedicated to processing attention. Importantly, feature coding in the early stage could predict the attentional effect in the late stage. Together, our results suggest an intricate interplay between visual feature and attention coding in the primate brain, which can be attributed to the differential functional connectivity and neural networks engaged in these processes.

## Introduction

Visual attention and visual object recognition are integral processes that govern how animals make sense of their visual environment. The primate brain efficiently processes and interprets the vast array of visual information encountered in their surroundings. Visual attention serves as the gateway to this process by enhancing the perception of relevant information while filtering out distractions, allowing the brain to selectively prioritize specific stimuli for further processing [1–3]. A substantial body of literature has documented the neural networks, pathways, and dynamics that dictate the brain’s selection, processing, and integration of information to achieve specific objectives [4, 5]. The complexity of visual attention involves a coordinated interaction among different brain regions and circuits, notably the prefrontal and temporal cortex [6–8]. These regions collaborate, playing a collective role in coordinating attentional resources and establishing intricate networks that dynamically adjust sensory processing in response to the goals and intentions of the animal [9].

In conjunction with visual attention, visual object recognition—the cognitive process responsible for identifying and categorizing objects in the visual field—involves integrating various visual features, such as shape, color, and texture, to create a coherent representation of an object. This intricate process relies on a sophisticated interplay of neural circuits, enabling the brain to identify and categorize objects based on a myriad of visual features [10]. Inferotemporal (IT) neurons play a critical role in the representation and analysis of visual objects [10–12]. In particular, IT neurons exhibit feature-based coding of objects, representing them across a broad and distributed population of neurons [13–16]. In a particular form of feature-based coding known as axis-based feature coding, IT neurons parametrically correlate with visual features along specific axes in feature space [17–20]. Neurons in downstream areas, such as the amygdala and hippocampus, likely receive this highly processed visual information as input and form high-level visual interpretations of stimuli [21]. Furthermore, IT neurons exhibit visually selective responses to natural stimuli even when they are embedded in complex natural scenes [22].

The interaction between visual attention and object processing is a dynamic and intricate dance within the brain. Specifically, visual attention and feature coding converge on the IT and V4 regions. The parallel processing of stimulus features guides visual search [23], and indeed feature attention predicts the efficiency of target detection [24]. While the analysis and encoding of visual features in V4 and IT neurons play a critical role in visual attention [8, 23–27], the precise nature of the interaction between visual attention and neural object coding remains unclear. This study systematically addressed this question by training macaques to perform a free-gaze visual search task using natural face and object stimuli, allowing for detailed analysis of visual features. We simultaneously recorded a large number of 10,639 units across multiple attention and visual coding areas, of which 3,931 units exhibited a focal foveal receptive field (RF). We hypothesized that neurons encoding visual attention and features overlapped in V4 and IT, suggesting an integration of these two processes. Importantly, we investigated whether neurons encoding visual features exhibited a different attentional response profile and functional connectivity compared to those not encoding visual features. Our investigation also systematically examined the modulation of attention on neuronal representational geometry and whether such modulation was more pronounced for neurons encoding visual features.

## Results

### Visual feature coding

Two monkeys performed a free-gaze visual search task, where their objective was to fixate on one of the two search targets that matched the same category as the cue (**Fig. 1A**). Both monkeys performed the task proficiently, with accuracy rates of 91.78%±0.19% for monkey S and 85.23%±0.41% for monkey E (see [28] for detailed behavioral and eye movement analyses). Notably, monkeys were trained to search for faces and houses that belonged to the same visual category as the cue but were different images, allowing us to investigate visual attention while utilizing a diverse range of images for studying visual object processing (**Fig. 1B**; see **Fig. S1** for characterization of the stimuli). Here, we focused on units with a focal foveal RF to facilitate the comparison between visual feature coding and attention coding, leading to 1624 units from area V4, 658 units from TE, 761 units from TEO, and 888 units from the orbitofrontal cortex (OFC) for further analysis (**Fig. 1C**; see **Fig. S2** for detailed characterization of the recording sites).

**Fig. 1.**
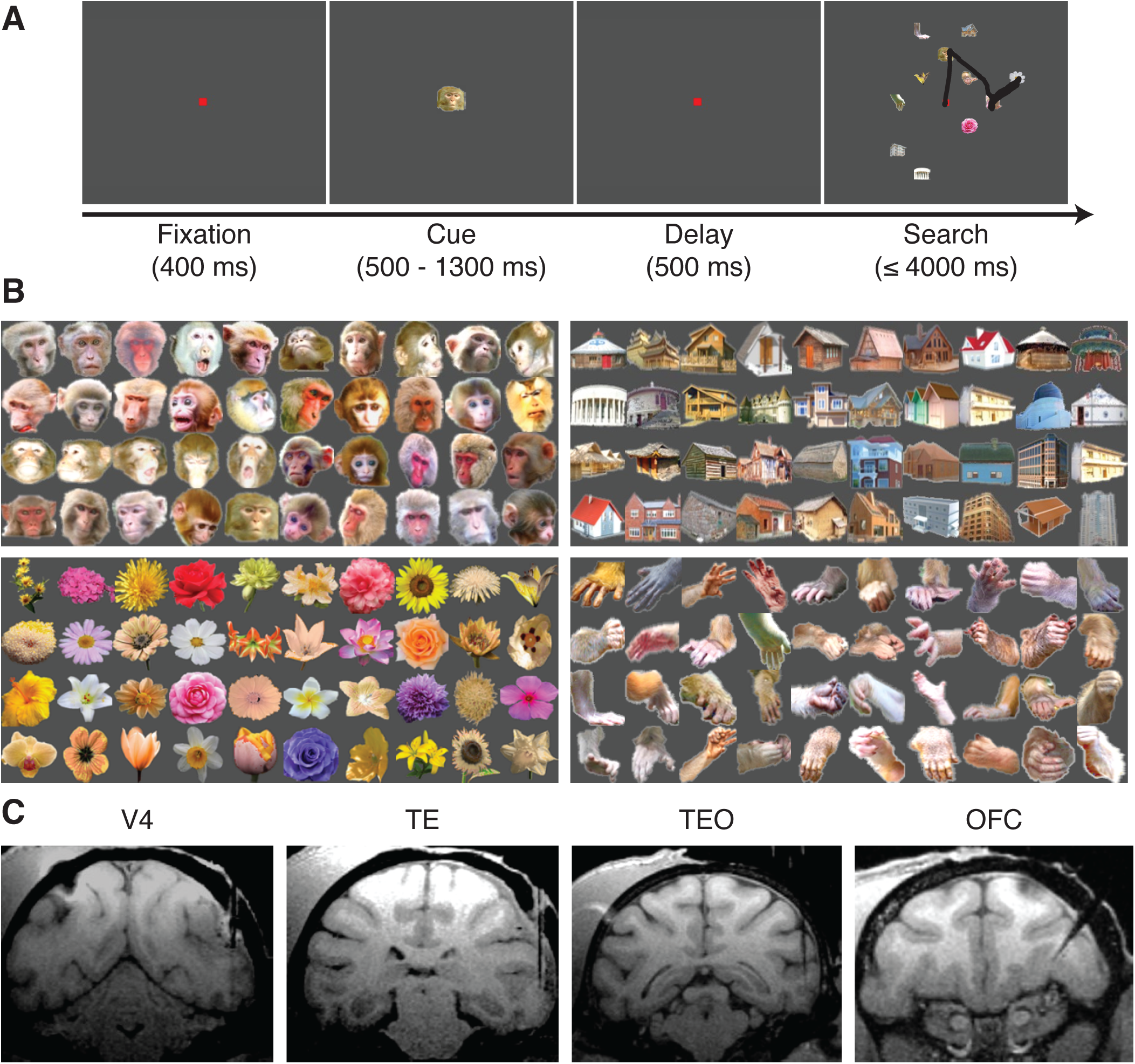
Task, stimuli, and recording sites. **(A)** Task. Monkeys initiated the trial by fixating on a central point for 400 ms. A cue was then presented for 500 to 1300 ms. After a delay of 500 ms, the search array with 11 items appeared. Monkeys were required to fixate on one of the two search targets that belonged to the same category as the cue for at least 800 ms to receive a juice reward. **(B)** Stimuli. Four categories of visual objects (40 images per category) were used for neural recordings. **(C)** MRI images show the typical recording regions of V4, TE, TEO, and the orbitofrontal cortex (OFC).

We first analyzed visual feature coding in each brain area using established approaches that are able to reveal whether a unit encodes a linear combination of deep neural network (DNN) features [19, 29] (see **Methods** for details). Using fixations on distractors that covered all stimulus items, we showed that IT (including both TE and TEO) and V4 units parametrically correlated with visual features along specific axes of the visual feature space (corresponding to a linear combination of DNN features; see **Fig. 2A-D** for examples and **Fig. 2E** for group summary), exhibiting an *axis code* for visual object representation. These units are referred to as *axis-coding units*; and this result is consistent with prior studies showing axis coding in the primate IT and V4 [17–20]. TE (31.76%; binomial P = 2.20×10^−15^; **Fig. 2E**) had the highest percentage of axis-coding units, which was significantly higher than that of TEO (11.96%; binomial P = 4.31×10^−14^; χ^2^-test: P < 10^−20^) and V4 (18.10%; binomial P = 6.65×10^−37^; χ^2^-test: P = 9.99×10^−13^); whereas OFC did not have an above-chance number of axis-coding units (3.15%). It is worth noting that we derived similar results when using DNN features from other layers (**Fig. S3A-D**), and our results also remained robust to features extracted using other DNNs (**Fig. S3E, F**).

**Fig. 2.**
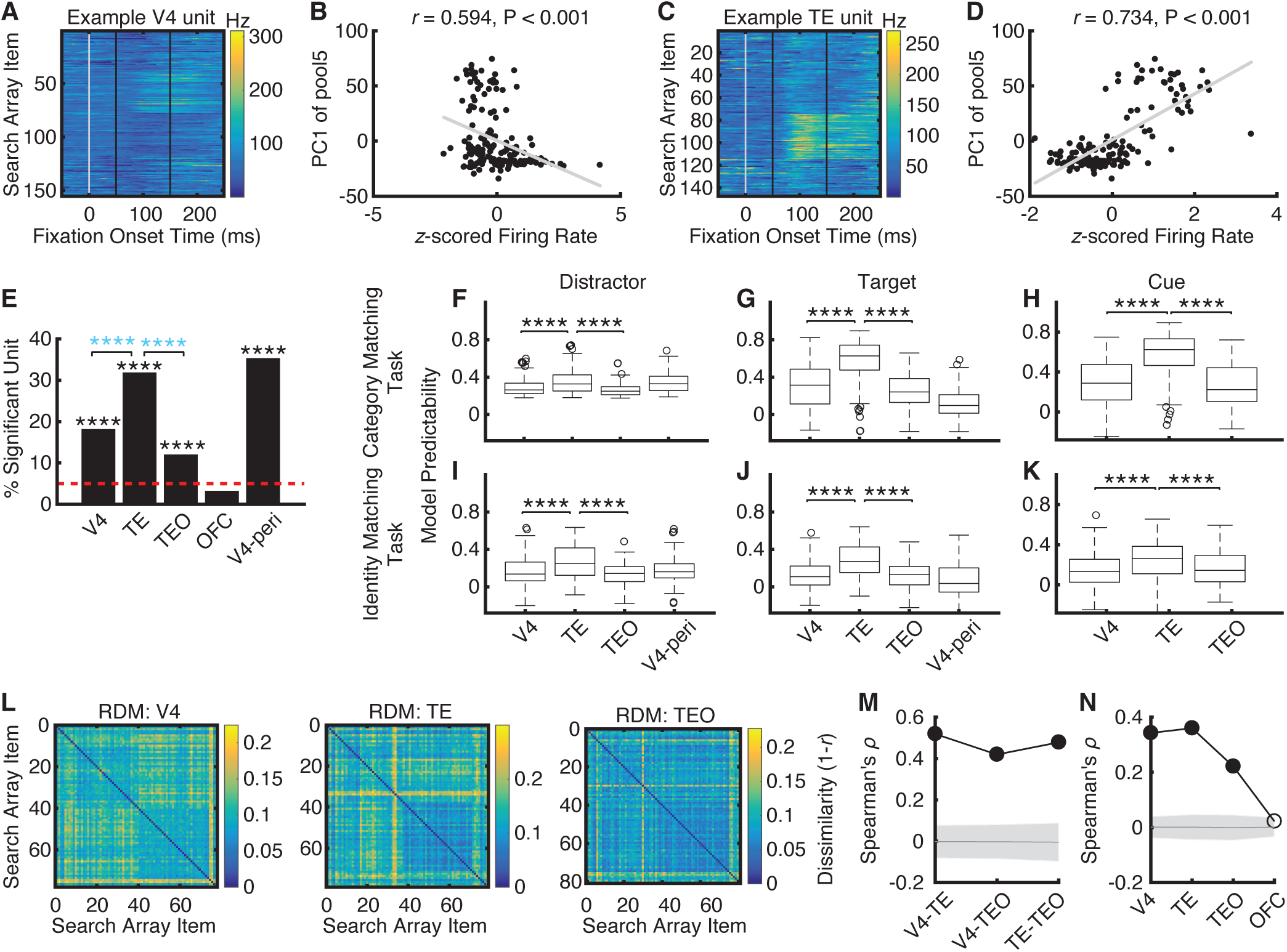
Axis-coding units. **(A-D)** Two example units showing axis-based feature coding. **(A, B)** A unit from V4. **(C, D)** A unit from TE. **(A, C)** Neural response to each search item. Time 0 denotes the fixation onset. Color coding shows the firing rate. **(B, D)** Correlation between the firing rate and the first principal component (PC1) of the feature map. Each dot represents a stimulus (i.e., search item), and the gray line denotes the linear fit. Note that both units had a significant relationship with the feature map (partial least squares [PLS] regression, permutation P < 0.001), and we show the correlation with PC1 for illustration purposes. **(E)** The proportion of units demonstrating axis-based feature coding for each brain area. Black asterisks indicate a significant above-chance (5%) number of units (binomial test). Blue asterisks indicate a significant difference between brain areas (χ^2^-test). ****: P < 0.0001. **(F-K)** Model predictability for each brain area. Predictability of PLS models was assessed using the Pearson correlation between the predicted and actual neural response in the test dataset. On each box, the central mark is the median across units, the edges of the box are the 25th and 75th percentiles, the whiskers extend to the most extreme data points the algorithm considers to be not outliers, and the circles denote the outliers. All box plots in this and subsequent figures follow the same convention. Asterisks indicate a significant difference between brain areas using two-tailed two-sample *t*-test. ****: P < 0.0001. **(F-H)** Category matching task. Monkeys were required to fixate on one of the two search targets that belonged to the same category as the cue (as shown in Fig. 1A). **(I-K)** Identity matching task. There was only one search target, and monkeys were required to fixate on this identical search target as the cue. **(F, I)** Analysis based on fixations on distractors. **(G, J)** Analysis based on fixations on targets. **(H, K)** Analysis based on fixations on cues. V4-peri: V4 units with a peripheral receptive field. **(L)** Representational dissimilarity matrices (RDMs) of search array items for each brain area. Color coding shows dissimilarity values (1−*r*). **(M)** Correlation between neural RDMs. The V4 RDM was significantly correlated with the TE RDM and TEO RDM, and the TE RDM was significantly correlated with the TEO RDM. **(N)** Correlation between neural RDM and deep neural network (DNN) feature RDM (VGG-16 layer pool5). Solid circles represent a significant correlation (permutation test: P < 0.05, Bonferroni-corrected across comparisons). Shaded area denotes ±SD across permutation runs.

The regression coefficients indicated the strength of axis coding (see **Methods**). We observed that TE exhibited the strongest axis coding compared to V4 and TEO (**Fig. 2F**; Ps < 0.0001), suggesting that the parametric encoding of complex visual features was most prominent in TE. Notably, this was also the case for fixations on targets (**Fig. 2G**) and fixations on cues (**Fig. 2H**). In addition, we further replicated this finding using a separate identity matching task where monkeys were required to search for an identical target as the cue (in contrast to the above category matching task, there was only one search target in the identity matching task). Specifically, TE exhibited the strongest axis coding compared to V4 and TEO for fixations on distractors (**Fig. 2I**), fixations on targets (**Fig. 2J**), and fixations on cues (**Fig. 2K**; all Ps < 0.0001). Furthermore, we found that V4 units with a focal peripheral RF (i.e., only one search item was encompassed by the RF) had similar visual feature coding as foveal units (**Fig. 2F, G, I, J**; note that TE and TEO did not have sufficient units with a focal peripheral RF for this analysis).

We next analyzed whether information flowed across brain areas using a representational similarity analysis (RSA) [30]. We found that the neuronal populations in V4, TE, and TEO (**Fig. 2L**) shared a similar representational structure (**Fig. 2M**; permutation Ps < 0.001), suggesting that information transitioned across the visual object processing areas. The brain areas exhibiting axis coding (i.e., V4, TE, and TEO; but not OFC) also demonstrated representational similarity with the DNN feature space (**Fig. 2N**). Lastly, we confirmed that axis coding correlated with visual category selectivity (**Fig. S4**).

Together, our results confirmed visual feature coding along the ventral processing pathway, which culminated in TE. Moreover, axis coding of visual features was present in different attentional contexts (distractors vs. targets vs. cue) and tasks (category matching vs. identity matching) and may transition across brain areas.

### Axis-coding units exhibit a different attentional effect

How are axis-coding units related to attention coding? We first identified *attention-selective units* that differentiated fixations on targets and distractors (**Methods**). We found that V4 (21.43%; binomial P = 6.65×10^−37^; **Fig. 3A**), TE (25.53%; binomial P = 2.20×10^−15^; **Fig. 3B**), and TEO (24.57%; binomial P = 1.12×10^−17^; **Fig. 3C**) all had an above-chance population of attention-selective units. Interestingly, axis-coding units were more likely to be attention-selective units in TE (i.e., the proportion of attention-selective units within axis-coding units [*n*_Both_ / *n*_Axis-Coding_] was significantly higher than the proportion of attention-selective units within all units [*n*_Attentive-Selective_ / *n*_All_]; χ^2^-test: P = 1.56×10^−3^) and V4 (P = 0.02). However, this was not the case for TEO units. Therefore, these results suggest that visual feature coding and attention coding interacted in TE and V4 but remained separate in TEO. We further correlated the effects of visual feature coding and attention, for all units from a brain area regardless of selectivity. We found that visual feature coding was correlated with attention in TE (*r* = 0.085, P = 0.03; **Fig. 3F**) but not in V4 (**Fig. 3E**) and TEO (**Fig. 3G**).

**Fig. 3.**
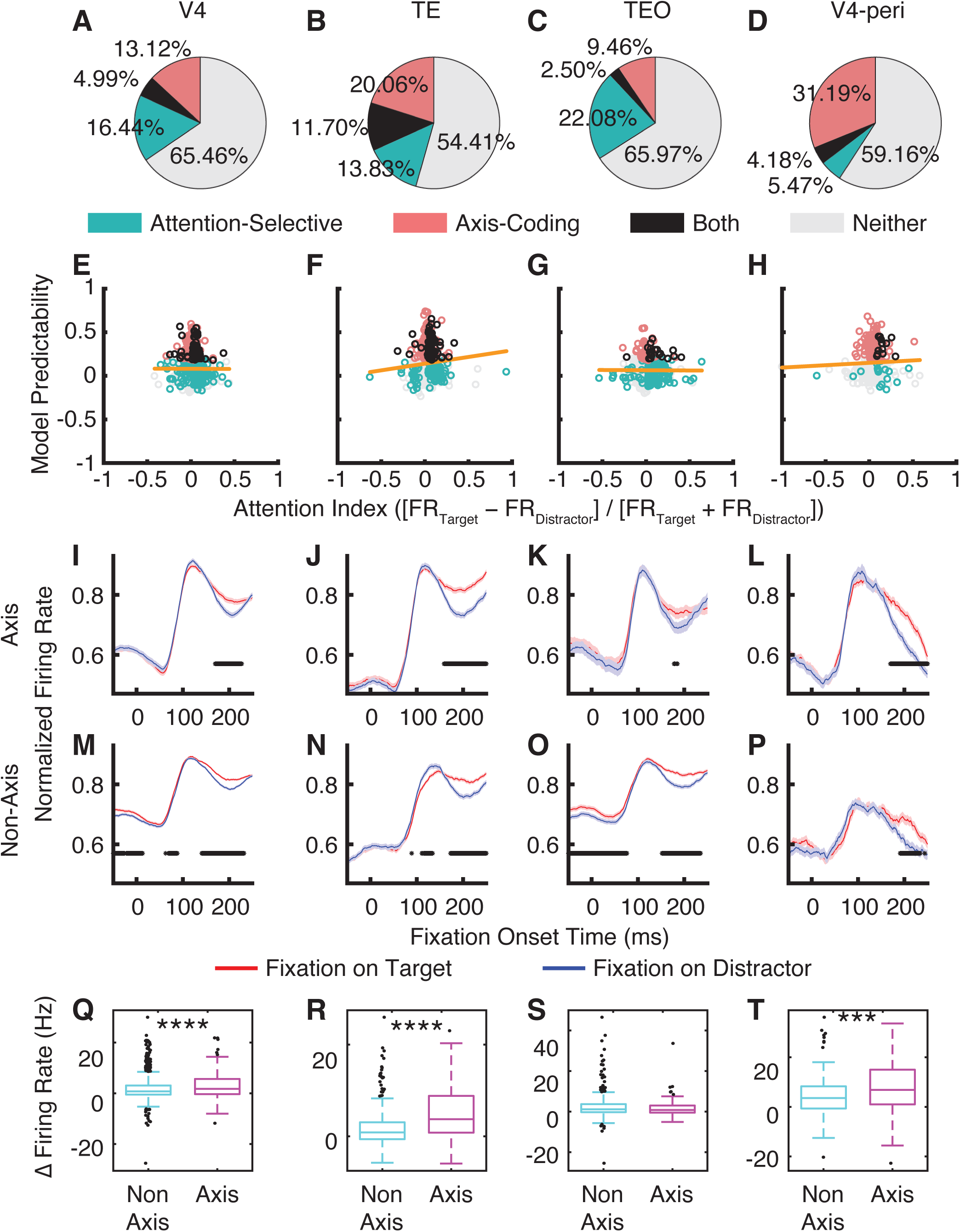
Attentional effect. **(A-D)** Population summary of attention-selective and axis-coding units. **(E-H)** Correlation between the strength of visual feature coding (shown by model predictability; see **Methods**) and the strength of attention coding ([FR_Target_ − FR_Distractor_] / [FR_Target_ + FR_Distractor_]). Each circle represents a unit. Color coding shows the unit type (green: attention-selective; red: axis-coding; black: both; gray: neither). The yellow line is the linear fit for all units. **(I-P)** Attentional effect. **(I-L)** Axis-coding units. **(M-P)** Non-axis-coding units. Shaded area denotes ±SEM across units. The black bars illustrate the time points with a significant difference between fixations on targets (red) and fixations on distractors (blue; two-tailed paired *t*-test, P < 0.05, Bonferroni-corrected across all time points). **(Q-T)** Comparison of attentional effect between axis-coding and non-axis-coding units for each brain area. Each box plot shows the difference in firing rate between fixations on targets and distractors. Asterisks indicate a significant difference using two-tailed two-sample *t*-test. ***: P < 0.001, and ****: P < 0.0001. **(A, E, I, M, Q)** V4. **(B, F, J, N, R)** TE. **(C, G, K, O, S)** TEO. **(D, H, L, P, T)** V4 units with a focal peripheral receptive field.

We next investigated whether units with versus without visual feature coding (i.e., axis coding) exhibited a differential response profile for attention. We first showed that across brain areas, both axis-coding (**Fig. 3I-K**) and non-axis-coding (**Fig. 3M-O**) units exhibited attentional effects during visual search. However, axis-coding units exhibited a stronger attentional effect than non-axis-coding units in V4 (**Fig. 3I, M, Q**; two-tailed two-sample *t*-test: *t*(1622) = 4.01, P = 6.32×10^−5^) and TE (**Fig. 3J, N, R**; *t*(656) = 8.74, P = 1.92×10^−17^) but not in TEO (**Fig. 3K, O, S**; *t*(759) = 0.63, P = 0.53). Therefore, attention coding interplayed with visual feature coding. It is important to note that axis-coding units were selected in an earlier time window (50-150 ms after fixation onset, when attentional effect barely emerged; **Fig. 2A, C**) than attentional effect (150-225 ms after fixation onset; **Fig. 3I-P**), suggesting that feature processing not only preceded attention but also predicted subsequent attentional effects (see also below).

Furthermore, while V4 units with a focal peripheral RF had a lower percentage of attention-selective units compared to units with a foveal RF (9.65%; binomial P = 6.18×10^−4^; **Fig. 3D**), the attentional effect was again stronger in axis-coding units compared to non-axis-coding units (**Fig. 3L, P, T**; *t*(309) = 3.88, P = 1.30×10^−4^), suggesting that visual feature coding and attention also interacted in the peripheral RF.

Together, we revealed units showing multiplexing functions for both attention and visual feature coding, with these two forms of coding primarily converged at TE. Importantly, units exhibiting visual feature coding had a stronger attentional effect in V4 and TE, suggesting an interaction between attention and visual feature coding.

### Attention modulates neural object representations

Above, we have demonstrated a differential response profile for axis-coding units when they encode visual attention. How does attention modulate neural object representations? Our previous work has shown that familiarity and familiarization modulate the population geometry of faces [31]. Utilizing this established approach, we quantified the population representational geometry of the units as a function of attentional contexts. It is worth noting that if all units change their response proportionally, the angle between the neuronal vectors will not change; otherwise, a change in the angle will suggest a change in the population geometry.

First, we showed that in axis-coding units, attention (i.e., fixations on targets compared to fixations on distractors) increased firing rate in the late time window (150 ms to 225 ms after fixation onset; **Fig. 4A**) in V4 (two-tailed paired *t*-test: *t*(63) = 8.20, P = 1.57×10^−11^) and TE (*t*(61) = 5.97, P = 1.29×10^−7^). Importantly, we found that both the neuronal distance (**Fig. 4E**; see **Fig. 4B-D** for illustration; V4: *t*(2015) = 28.93, P = 3.34×10^−154^; TE: *t*(1890) = 35.46, P = 1.56×10^−211^; TEO: *t*(2015) = 15.52, P = 2.16×10^−51^) and the angle between the neuronal vectors (**Fig. 4F**; V4: *t*(2015) = 14.95, P = 5.15×10^−48^; TEO: *t*(2015) = 15.80, P = 3.92×10^−53^; except TE: *t*(1890) = 1.40, P = 0.16) *increased* for targets compared to distractors, suggesting that neural representations of targets became more distinct, which could in turn facilitate target detection. Notably, such enhancement in neuronal representational distance and angle even happened in TEO where firing rate did not increase for search targets (**Fig. 4A**; *t*(63) = −1.96, P = 0.054), suggesting that changes in neuronal representational geometry could be dissociated from changes in firing rate.

**Fig. 4.**
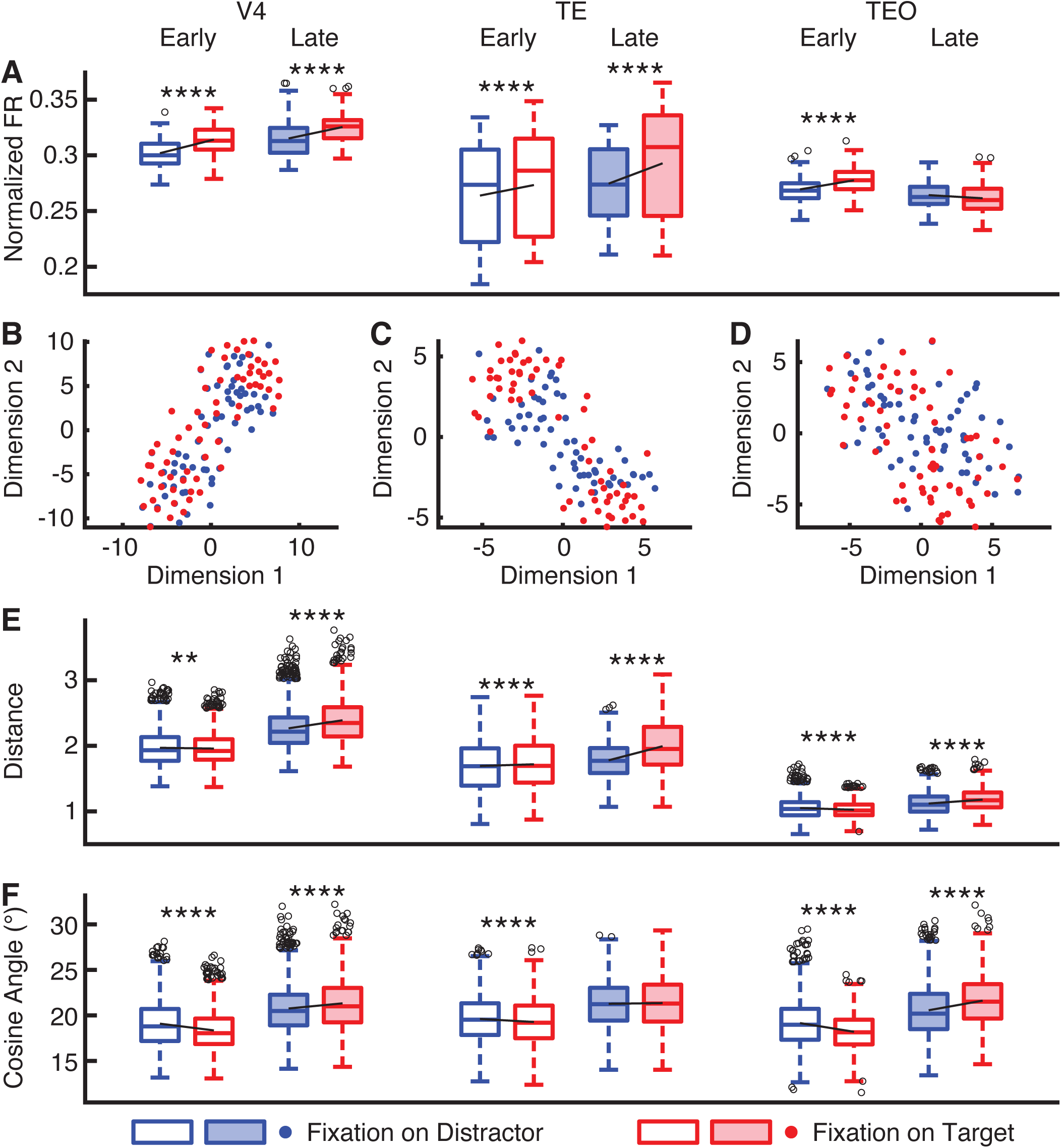
Attention modulation of neuronal representational geometry. **(A)** Normalized firing rate (FR). **(B-D)** Distribution of stimuli in the neuronal feature space (constructed using principal component analysis [PCA]). Each dot represents a stimulus. **(B)** V4. **(C)** TE. **(D)** TEO. **(E)** Representational distance for the population of neurons. **(F)** Angle between the neuronal vectors. Open boxes: early time window (50 ms to 150 ms after fixation onset). Solid boxes: late time window (150 ms to 225 ms after fixation onset). Asterisks indicate a significant difference between conditions (distractor vs. target) using two-tailed paired *t*-test. *: P < 0.05, **: P < 0.01, ***: P < 0.001, and ****: P < 0.0001.

Second, we found that in the early time window (50 ms to 150 ms after fixation onset), even though firing rate increased for fixations on targets across brain areas (**Fig. 4A**; V4: *t*(63) = 11.77, P = 1.49×10^−17^; TE: *t*(61) = 6.30, P = 3.58×10^−8^; TEO: *t*(63) = 6.89, P = 3.05×10^−9^), attention did *not enhance* the neuronal distance (**Fig. 4E**; V4: *t*(2015) = −3.15, P = 0.0017; TEO: *t*(2015) = −7.73, P = 1.71×10^−14^; except TE: *t*(1890) = 6.26, P = 4.76×10^−10^) or the angle between the neuronal vectors (**Fig. 4F**; V4: *t*(2015) = −21.92, P = 1.14×10^−95^; TE: *t*(1890) = −7.48, P = 1.13×10^−13^; TEO: *t*(2015) = −16.06, P = 1.12×10^−54^), suggesting that attention modulated neuronal representations in a later stage.

Third, we examined whether attention modulation of neuronal representations was particularly pronounced for axis-coding units compared to non-axis-coding units (**Fig. 5**). Indeed, in the late time window, axis-coding units across brain areas exhibited a stronger attention modulation for both the neuronal distance (**Fig. 5B**; V4: *t*(3667) = 29.07, P = 2.40×10^−167^; TE: *t*(3719) = 13.90, P = 7.69×10^−43^; TEO: *t*(3725) = 10.16, P = 5.87×10^−24^) and the angle between the neuronal vectors (**Fig. 5C**; TE: *t*(3719) = 7.77, P = 1.00×10^−14^; TEO: *t*(3725) = 8.77, P = 2.77×10^−18^; except V4: *t*(3667) = 1.04, P = 0.30), showing a disproportionate attention modulation for units encoding visual features. It is worth noting that changes in neuronal representations were greater for axis-coding units even if changes in firing rate were greater (**Fig. 5A**; V4: *t*(120) = 14.79, P = 8.02×10^−29^), smaller (TE: *t*(121) = −2.12, P = 3.58×10^−2^), or similar (TEO: *t*(121) = 0.05, P = 0.96) compared to non-axis-coding units, again indicating a dissociation between firing rate and neuronal representational geometry.

**Fig. 5.**
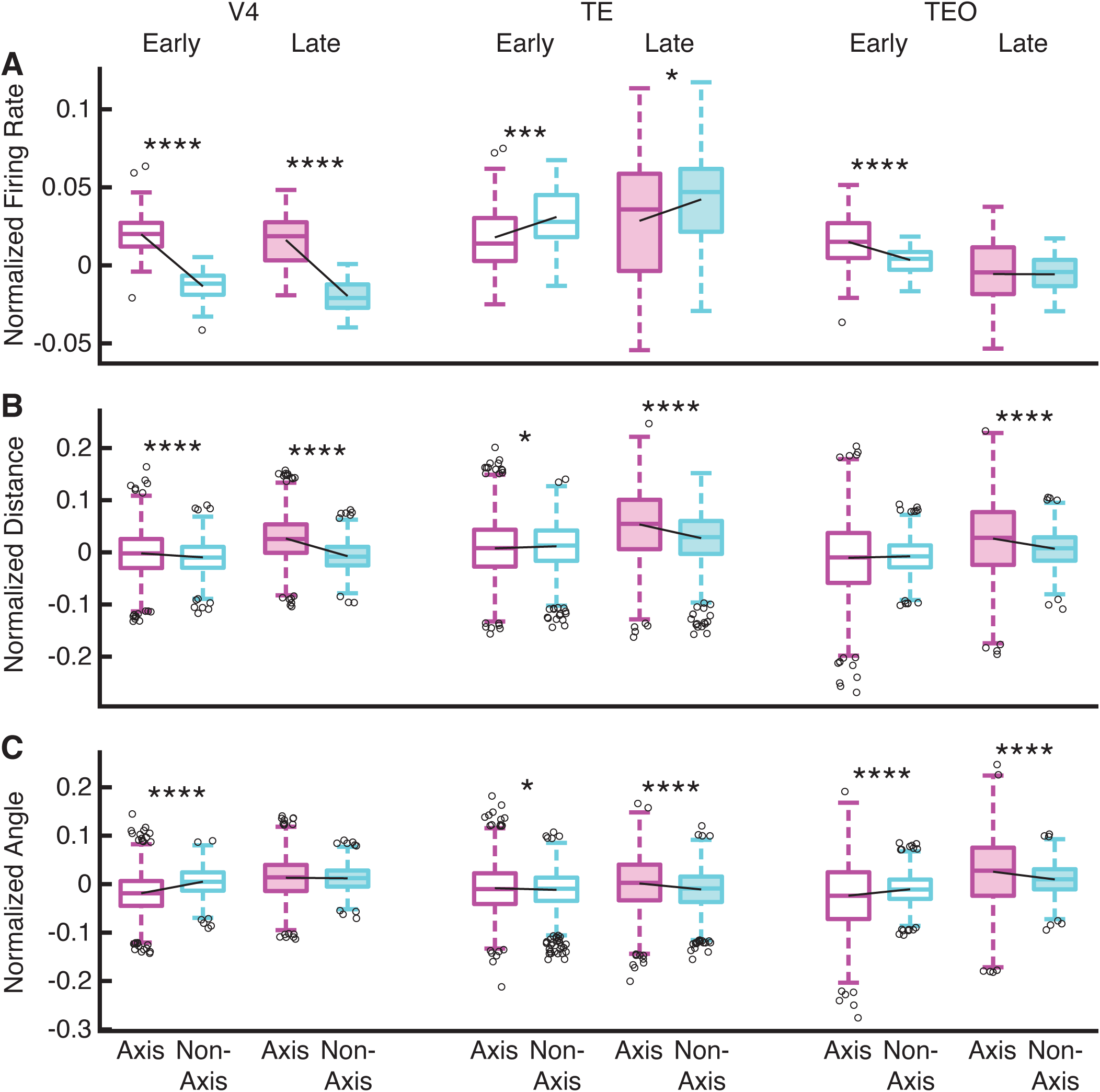
Differential attention modulation of neuronal representational geometry between axis-coding and non-axis-coding units. **(A)** Normalized firing rate ([FR_Target_ − FR_Distractor_] / [FR_Target_ + FR_Distractor_]). **(B)** Normalized representational distance for the population of neurons ([Distance_Target_ − Distance_Distractor_] / [Distance_Target_ + Distance_Distractor_]). **(C)** Normalized angle between the neuronal vectors ([Angle_Target_ − Angle_Distractor_] / [Angle_Target_ + Angle_Distractor_]). Open boxes: early time window (50 ms to 150 ms after fixation onset). Solid boxes: late time window (150 ms to 225 ms after fixation onset). Asterisks indicate a significant difference between axis-coding and non-axis-coding units using two-tailed two-sample *t*-test. *: P < 0.05, **: P < 0.01, ***: P < 0.001, and ****: P < 0.0001.

Lastly, in the early time window, although changes in firing rate were significantly different between axis-coding and non-axis-coding units (**Fig. 5A**; all Ps < 0.001), we did not observe a consistent pattern of differences in neuronal representational geometry (**Fig. 5B, C**).

Together, we demonstrated that attention to search targets enhanced pattern separation of the stimuli across brain areas, and such enhancement was more pronounced for units encoding visual features.

### Disproportionate target-induced desynchronization for axis-coding versus non-axis-coding units

Are axis-coding and non-axis-coding units engaged in the same functional network? To answer this question, we analyzed the coherence between spikes and local field potentials (LFPs) recorded simultaneously across brain areas (see **Methods**). We included spikes from V4 and IT (OFC and lateral prefrontal cortex [LPFC] were excluded due to having fewer than 20 axis-coding units) and LFPs from all four brain areas. We found that for both axis-coding and non-axis-coding units, spikes desynchronized with LFPs in the theta frequency band for fixations on targets compared to fixations on distractors (**Fig. 6**). Importantly, axis-coding units demonstrated a stronger target-induced desynchronization between V4 spike and V4 LFP (**Fig. 6A**; two-tailed two-sample *t*-test: *t*(29404) = 2.98, P = 0.00288), between V4 spike and IT LFP (**Fig. 6B**; *t*(7801) = 2.06, P = 0.0395), between V4 spike and OFC LFP (**Fig. 6C**; *t*(4500) = 2.03, P = 0.0424), between IT spike and V4 LFP (**Fig. 6E**; *t*(7979) = 10.27, P = 1.42×10^−24^), between IT spike and IT LFP (**Fig. 6F**; *t*(22658) = 9.36, P = 8.77×10^−21^), and between IT spike and OFC LFP (**Fig. 6G**; *t*(5037) = 4.62, P = 4.01×10^−6^), suggesting that axis-coding units disproportionately engaged the attention network compared to non-axis-coding units. The systematic differences between axis-coding and non-axis-coding units also indicated differential top-down modulation from the prefrontal cortex.

**Fig. 6.**
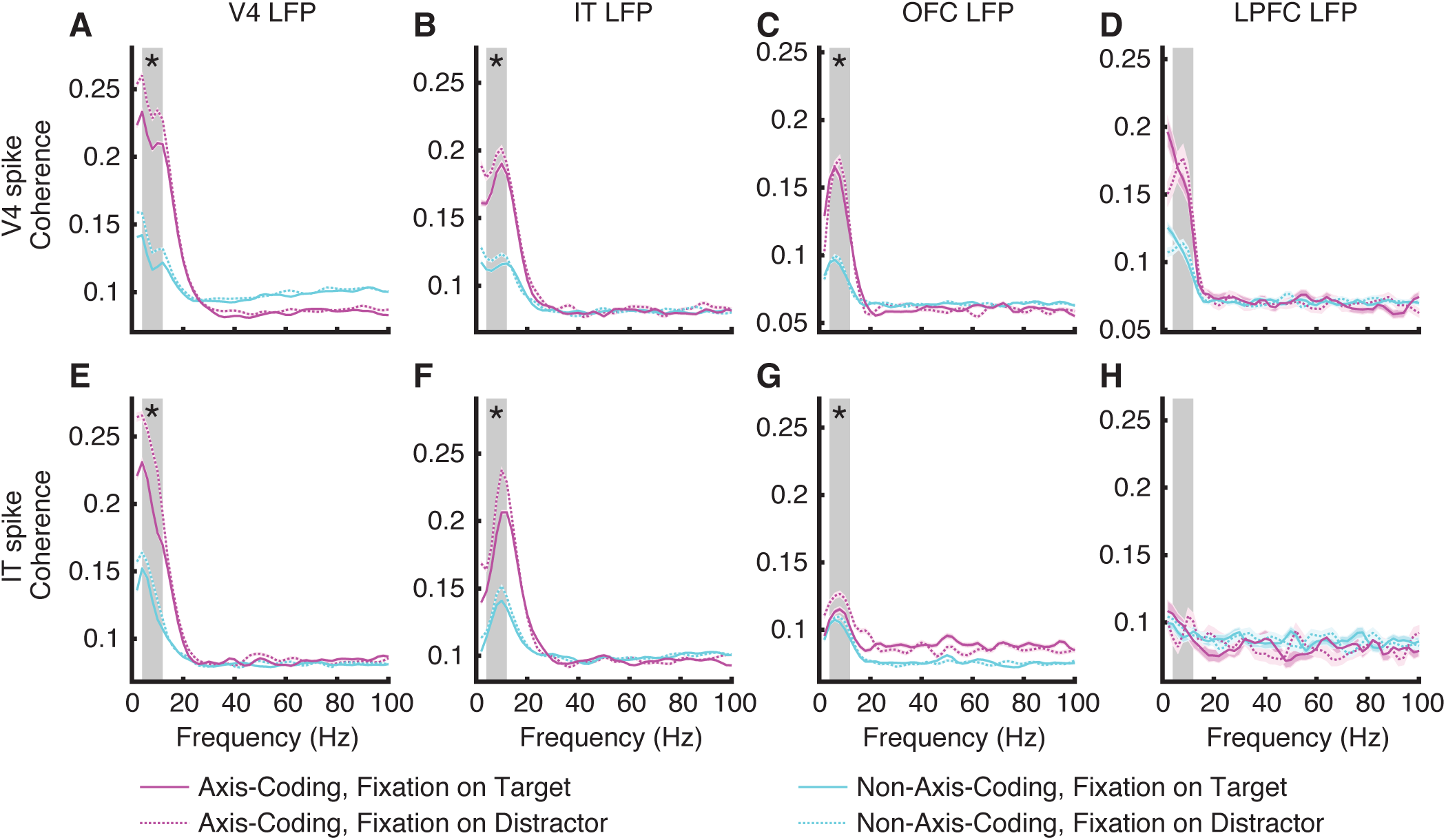
Spike-LFP coherence. **(A)** V4 spike-V4 LFP. **(B)** V4 spike-IT LFP. **(C)** V4 spike-OFC LFP. **(D)** V4 spike-LPFC LFP. **(E)** IT spike-V4 LFP. **(F)** IT spike-IT LFP. **(G)** IT spike-OFC LFP. **(H)** IT spike-LPFC LFP. Magenta: axis-coding units. Cyan: non-axis-coding units. Solid line: fixation on targets. Dotted line: fixation on distractors. Magenta and cyan shaded areas denote ±SEM across spike-LFP pairs. Gray shaded area denotes the theta frequency band (4 - 12 Hz). Asterisks indicate a significant difference in target-induced desynchronization (i.e., the reduction in spike-LFP coherence for fixations on targets compared to fixations on distractors, averaged across the theta frequency band) between axis-coding units and non-axis-coding units using two-tailed two-sample *t*-test.

### Directional theta influence across brain areas for axis-coding versus non-axis-coding units

To investigate the direction of interactions between brain areas, we performed a Granger causality analysis based on spikes and LFPs in the theta frequency band (see **Fig. S5** for analyses across frequencies). We first analyzed the influence of spikes on LFPs (**Fig. 7A, C, D**; **Fig. S5A-H**). We found that attention modulated interactions between brain areas when comparing fixations on targets versus distractors, with targets inducing a decrease in Granger causality (i.e., desynchronization). Importantly, such modulation was more pronounced for axis-coding versus non-axis-coding units (**Fig. 7C, D**). Specifically, the influence of V4 spike on V4 LFP (**Fig. S5A**; two-tailed two-sample *t*-test: *t*(29404) = 3.54, P = 4.01×10^−4^), the influence of V4 spike on LPFC LFP (**Fig. 7A**; **Fig. S5D**; *t*(689) = 3.62, P = 3.19×10^−4^), and the influence of IT spike on IT LFP (**Fig. S5F**; *t*(22658) = 2.46, P = 0.0138) were more strongly modulated in axis-coding units compared to non-axis-coding units.

**Fig. 7.**
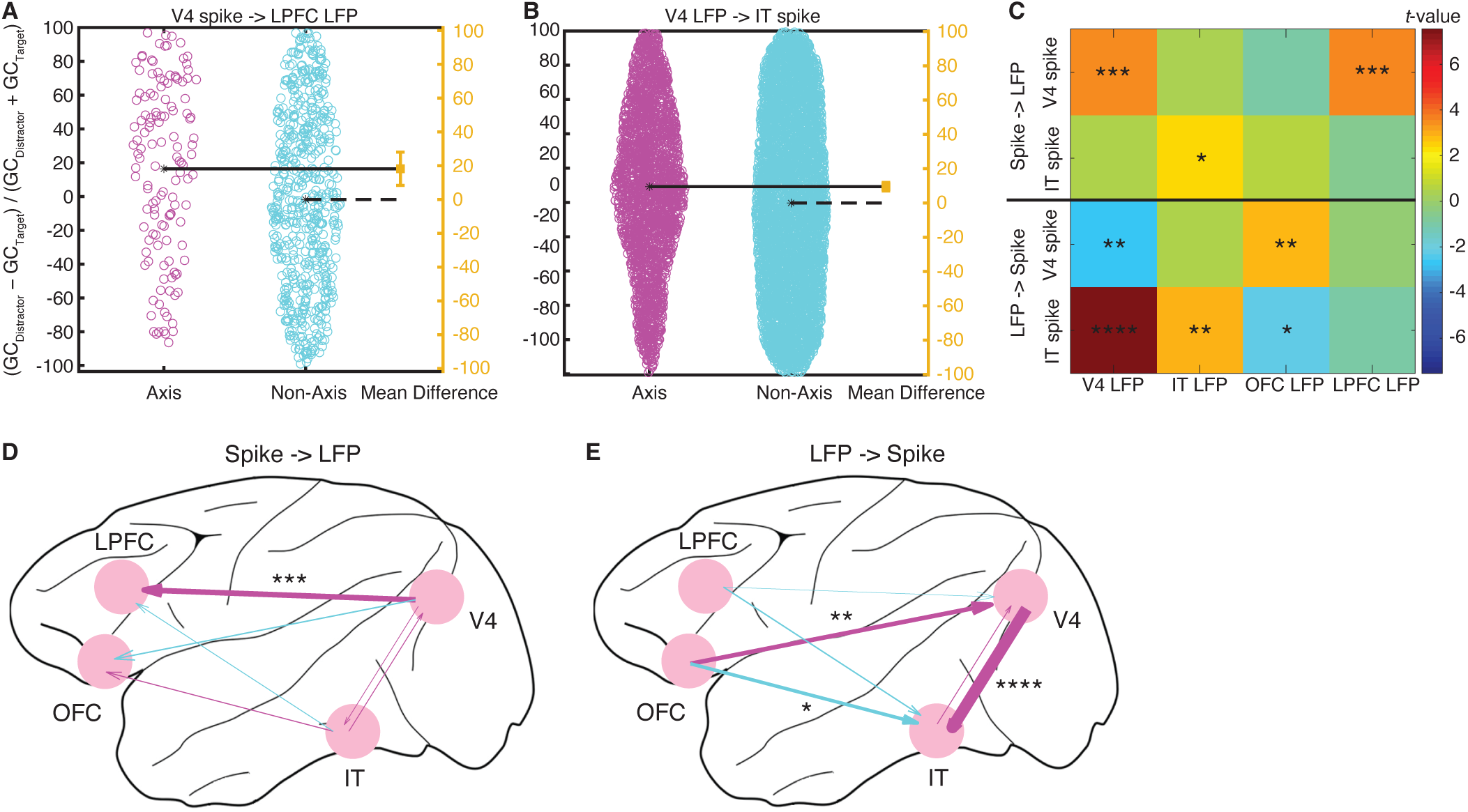
Granger causality (GC). **(A, B)** Compared to non-axis-coding units, axis-coding units had a stronger reduction in Granger causality between brain areas ([GC_Distractor_ − GC_Target_] / [GC_Distractor_ + GC_Target_]). Each circle represents a spike-LFP pair. The mean of the non-axis-coding units corresponds to the zero effect size and the mean of the axis-coding units corresponds to the value of the effect size on the effect size axis (yellow). The vertical error bar in yellow indicates the actual mean-difference effect size value and the confidence intervals. **(A)** V4 spike influence on LPFC LFP. **(B)** V4 LFP influence on IT spike. **(C)** Summary of Granger causality for each spike-LFP pair. Color coding shows the *t*-values of the two-tailed two-sample *t*-test (axis − non-axis). Asterisks indicate a significant difference between axis-coding units and non-axis-coding units using two-tailed two-sample *t*-test. *: P < 0.05, **: P < 0.01, ***: P < 0.001, and ****: P < 0.0001. **(D, E)** Differences in target-induced reduction of cross-area Granger causality between axis-coding and non-axis-coding units. The thickness of the arrow is proportional to the *t*-value of the two-tailed two-sample *t*-test. Red: axis-coding > non-axis-coding. Blue: non-axis-coding > axis-coding. **(D)** Spike influence on LFP. **(E)** LFP influence on spike.

We also analyzed the influence of LFPs on spikes (**Fig. 7C, E**; **Fig. S5I-P**). Again, we found that attention modulated interactions between brain areas, and such modulation was disproportionate for axis-coding units (**Fig. 7C, E**). Notably, the influence of OFC LFP on V4 spike (**Fig. S5K**; *t*(4500) = 2.91, P = 0.00367), the influence of V4 LFP on IT spike (**Fig. 7B**; **Fig. S5M**; *t*(7979) = 7.54, P = 5.05×10^−14^), and the influence of IT LFP on IT spike (**Fig. S5N**; *t*(22658) = 2.98, P = 0.00292) were more strongly modulated in axis-coding units; whereas the influence of V4 LFP on V4 spike (**Fig. S5I**; *t*(29404) = 2.60, P = 0.00930) and the influence of OFC LFP on IT spike (**Fig. S5O**; *t*(5037) = 2.19, P = 0.0285) were more strongly modulated in non-axis-coding units.

Together, our results reveal bidirectional influences between spikes and LFPs across brain areas. In particular, attention modulation of these influences is more pronounced for axis-coding units compared to non-axis-coding units, suggesting that axis-coding units differentially engage the attention neural network.

## Discussion

In this study, we recorded from a large number of units across attention and visual coding areas and demonstrated that units exhibiting visual feature coding (i.e., axis-coding units) displayed a distinct attentional response profile compared to non-axis-coding units. Moreover, axis-coding units exhibited different functional connectivity patterns to other brain areas when encoding attention, suggesting differential engagement with the attention neural network. Importantly, our results suggest two stages of processing. In the early stage (e.g., 50 ms to 150 ms after fixation onset), neurons primarily process visual features and exhibit axis coding. In the late stage (e.g., 150 ms to 225 ms after fixation onset), neurons primarily process attention. Notably, feature coding in the early stage could predict the attentional effect in the late stage. Together, our results suggest an intricate interplay between visual feature coding and attention coding in the primate brain, which can be attributed to the differential functional connectivity and neural networks engaged in these processes..

We trained macaques to perform a free-gaze visual search task using natural face and object stimuli, enabling detailed analysis of visual features. Our study is thus uniquely positioned to investigate the intricate interplay between attention coding and visual feature coding. Indeed, we identified units encoding both visual features and attention, resembling the multidimensional processing observed in the primate amygdala, where the same neurons encode valence, arousal, and visual features [32]. Neurons encoding visual features may form the basis of feature-based attention. Consistent with our present finding, it has been shown using a similar naturalistic free-gaze visual search task that V4 neurons not only exhibit visually driven response to features but also top-down modulation of response to search targets [26]. Notably, we found that units encoding visual features exhibited a different attentional effect and functional connectivity. Similar to our present findings, units in the human amygdala and hippocampus not only encode a visual attentional effect towards search targets but also encode visual categories [33]; however, in contrast, the units encoding attention and visual categories appear to be independent. This raises the question of whether and where these two processes diverge along the processing stream, a question that future analyses should address.

The location-independent property of feature-based attention makes it particularly well suited to selectively modify the neural representations of stimuli or parts within complex visual scenes that match the currently attended feature [34]. In this study, we found attention modulation of neural representational geometry, supporting a recent hypothesis that attention improves performance by reshaping stimulus representations to align with the readout [35]. Our present results can be interpreted in the framework of pattern separation [36, 37], the process of transforming similar representations or memories into dissimilar, non-overlapping representations, and are in line with the tuning sharpening in the primate IT cortex [38, 39]. Importantly, we found that the representational geometry changed differently in axis-coding versus non-axis-coding units, and we revealed the temporal dynamics (and specificity) for visual feature coding and attention modulation. Furthermore, it is worth noting that while neuronal representational distance and angle were calculated based on firing rate, changes in these measures could be dissociated from changes in firing rate (**Fig. 4** and **Fig. 5**).

We observed a desynchronization for attended stimuli (fixations on search targets) in the theta frequency band, as opposed to the same stimuli when they were unattended (fixations on distractors) in both V4 and IT. This finding aligns with previous research demonstrating desynchronization for attended stimuli in V4 within a similar frequency band [40, 41]. Desynchronization has been observed in instances of feature-based attention, where discrimination between target and distractor in the peripheral RF occurs, as well as during saccade selection, involving directing attention into (attention in) or out of (attention out) the peripheral RF. This pattern was evident in V4 spike-V4 LFP coherence, V4 spike-frontal eye field (FEF) LFP coherence, and FEF spike-V4 LFP coherence [41]. Furthermore, spike-field Granger causality can be used to reveal the modulatory effects that are inaccessible by traditional methods, such that spike->LFP Granger causality is modulated by the behavioral task, whereas LFP->spike Granger causality is mainly related to the average synaptic input [42]. Notably, in this study, we observed bidirectional differences in Granger causality between axis-coding and non-axis-coding units.

Our present result is consistent with top-down modulation from the frontal cortex to the temporal lobe [33, 43]. It has been shown that attention to faces versus houses enhanced the sensory responses in the fusiform face area (FFA) and parahippocampal place area (PPA), respectively. The increases in sensory responses are accompanied by induced gamma synchrony between the inferior frontal junction (IFJ) and either FFA or PPA, depending on which object is attended; and the IFJ directs the flow of visual processing during object-based attention, at least in part through coupled oscillations with specialized areas such as FFA and PPA [44]. In addition, individual PFC units synchronized to the LFP ensemble corresponding to the current task goal or rule and the neural ensemble encoding the behaviorally dominant rule exhibits increased ‘‘alpha’’ (6 - 16 Hz) synchrony when preparing to apply the alternative (weaker) rule [45], consistent with our prior report showing that different search goals (social vs. non-social) differentially engage the PFC. Our present result is also consistent with the rhythmic theory of attention that both perceptual sensitivity during covert spatial attention and the probability of overt exploratory movements are tethered to theta-band activity in the attention network [46].

In conclusion, our study sheds light on the complex relationship between visual feature coding and attention coding in the primate brain. We have demonstrated that units exhibiting visual feature coding display distinct attentional response profiles and functional connectivity compared to units not exhibiting feature coding, indicating a nuanced interplay between these cognitive processes. Future research exploring how these findings translate to behavioral outcomes and cognitive functions in both primates and humans could offer valuable implications for understanding attentional processes and cognitive control in diverse contexts.

## Acknowledgements

This research was supported by the NSF (BCS-1945230), NIH (R01MH129426), and AFOSR (FA9550-21-1-0088). The funders had no role in study design, data collection and analysis, decision to publish, or preparation of the manuscript.

## Author Contributions

J.Z. and H.Z. designed research. J.Z. and X.Z. performed experiments. J.Z., R.C., and S.W. analyzed data. J.Z., H.Z., and S.W. wrote the paper. All authors discussed the results and contributed toward the manuscript.

## Methods

### Subjects

Two male rhesus macaques, weighing 12 and 15 kg, were used in the study. The monkeys were implanted under aseptic conditions with a post to fix the head and recording chambers over areas V4, inferotemporal (IT) cortex (including both TE and TEO), lateral prefrontal cortex (LPFC), and orbitofrontal cortex (OFC). The localization of the chambers was based on MRI scans obtained before surgery. All experiments were performed with the approval of the Institutional Animal Care and Use Committee of Shenzhen Institutes of Advanced Technology, Chinese Academy of Sciences (No. SIAT-IRB-160223-NS-ZHH-A0187-003).

### Tasks and stimuli

Monkeys were trained to perform a free-gaze visual search task. A central fixation was presented for 400 ms, followed by a cue lasting 500 to 1300 ms. After a delay of 500 ms, the search array was on. The search array contained 11 items, including two targets, randomly selected from a total of 20 predefined locations. Monkeys were required to find either one of the two targets within 4000 ms and maintain fixation on the target for 800 ms to receive a juice reward. No constraints were placed on their search behavior to allow animals to perform the search naturally. Before the onset of the search array, monkeys were required to maintain a central fixation. The two target stimuli belonged to the same category as the cue stimulus, though they were distinct images. We utilized four categories of stimuli—face, house, flower, and hand—each comprising 40 images. The cue stimulus was randomly selected from the house or face stimuli with equal probability. The remaining 9 stimuli in the search array were drawn from the other three categories. Each stimulus subtended an area of approximately 2° × 2°, with the hue, saturation in the HSV color space, aspect ratio, and luminance of these images matched across categories. The 20 locations, covering the visual field of eccentricities from 5° to 11°, included 18 locations located symmetrically in the left and right visual field, with 9 on each side, and 2 locations on the vertical middle line.

A visually guided saccade task was employed to map the peripheral receptive fields (RFs) of recorded units. Following a 400-ms central fixation, a stimulus randomly appeared in one of the 20 locations, and monkeys were required to make a saccade to the stimulus within 500 ms and maintain fixation on it for 300 ms to receive a reward.

Behavioral experiments were conducted using the MonkeyLogic software (University of Chicago, IL), which presented the stimuli, monitored eye movements, and triggered the delivery of the reward.

### Electrophysiology

Single-unit and multi-unit spikes were recorded from V4, IT, LPFC, and OFC using 24- or 32-contact electrodes (V-Probe or S-Probe, Plexon Inc, Dallas, USA) in a 128-channel Cerebus System (Blackrock Microsystems, Salt Lake City, UT, USA). In most sessions, we recorded activities in two of the areas simultaneously. Neural signals were filtered between 250 Hz and 5 kHz, amplified, and digitized at 30 kHz to obtain spike data. The recording locations in V4, IT, LPFC, and OFC were verified with MRI. Eye movements were recorded using an infrared eye-tracking system (iViewX Hi-Speed, SensoMotoric Instruments (SMI), Teltow, Germany) at a sampling rate of 500 Hz.

### Data analysis: spike rate

Measurements of neural activity were obtained from spike density functions, which were generated by convolving the time of action potentials with a function that projects activity forward in time (Growth = 1 ms, Decay = 20 ms) and approximates an EPSP [47]. Spike rate of each unit was normalized by its maximal spike rate across conditions.

### Data analysis: receptive field

The visual response to the cue and the search array in the free-gaze visual search task was assessed by comparing the firing rate during the post-stimulus period (50 to 200 ms after cue/array onset) to the corresponding baseline (−150 to 0 ms relative to cue/array onset) using a Wilcoxon rank-sum test. Based on these responses, we classified units into three categories of RFs:

i. Units with a focal foveal RF: These units responded solely to the cue in the foveal region (P < 0.05) but not to the search array that included items in the periphery (P > 0.05).
ii. Units with a broad foveal RF: These units responded to both the cue and the search array.
iii. Units with a peripheral RF: These units only responded to the search array (P < 0.05) but not to the cue (P > 0.05). The RFs of these units were additionally mapped based on their activities in the visually guided saccade task.

Units not classified into one of the three categories were excluded from further analysis. In this study, our focus was on units with a focal foveal RF (category (i)) because we aimed to analyze the visual feature coding properties and compare fixations on targets versus distractors. We also analyzed the response in the V4 peripheral RF because V4 (but not TE or TEO) had a focal peripheral RF (i.e., only one search item was encompassed by the RF).

### Data analysis: category selectivity

We determined the category selectivity of each unit by comparing the response to face cues versus house cues in a time window of 50 to 200 ms after cue onset (Wilcoxon rank-sum test, P < 0.05). We further imposed a second criterion using a selectivity index similar to indices employed in previous IT studies [48, 49]. For each unit with a foveal RF, the response to face stimuli (*R*_face_) or house stimuli (*R*_house_) was calculated using the visual search task by subtracting the mean baseline activity (−150 to 0 ms relative to the onset of the cue) from the mean response to the face or house cue (50 to 200 ms after the onset of the cue). For each unit with a peripheral RF, *R*_face_ and *R*_house_ were calculated using the visually guided saccade task by subtracting the mean baseline activity (−150 to 0 ms relative to the peripheral stimulus onset) from the mean response to the saccade target (50 to 200 ms after the onset of the saccade target). The selectivity index (SI) was then defined as (*R*_face_ − *R*_house_) / (*R*_face_ + *R*_house_). SI was set to 1 when *R*_face_ > 0 and *R*_house_ < 0, and to −1 when *R*_face_ < 0 and *R*_house_ > 0. Face-selective units were required to have an *R*_face_ at least 130% of *R*_house_ (i.e., the corresponding SI was greater than 0.13). Similarly, house-selective units were required to have an *R*_house_ at least 130% of *R*_face_ (i.e., the corresponding SI was smaller than −0.13). Units were labeled as non-category-selective if the response to face cues versus house cues was not significantly different (P > 0.05). The remaining units that did not fit into any of the aforementioned types were classified as undefined units (i.e., there was a significant difference but did not meet the second criterion).

### Data analysis: selection of axis-coding units

We used the well-known deep neural network (DNN) implementation based on the VGG-16 convolutional neural network (CNN) architecture [50] to extract features for each image (replicated by AlexNet [51] and ResNet [52]; **Fig. S3**). Following the same procedure [53], fine-tuning of the top layer of each DNN was performed to confirm that the pre-trained model was able to discriminate the objects and ensure that the pre-trained model was suitable as a feature extractor. We also used the fine-tuning accuracy to determine the most suitable model for feature extraction.

To identify axis-coding units (i.e., units encoding a linear combination of visual features), we employed a partial least squares (PLS) regression with DNN feature maps. We used the mean firing rate in a time window of 50 ms to 150 ms after fixation onset as the response to each fixation [54, 55] (see also **Fig. 2A, C** for reference). The PLS method has been shown to be effective to study the neural response to DNN features [19, 29]. We used 4 components for each layer, explaining at least 60% of variance. We used a permutation test with 1000 runs to determine whether a unit encoded *a significant axis-coding model* (i.e., the unit encoded the dimensions of the feature space, demonstrating *axis coding*). In each run, we randomly shuffled the object labels and used 50% of the objects as the training dataset. We used the training dataset to construct a model (i.e., deriving regression coefficients), predicted responses using this model for each object in the remaining 50% of objects (i.e., test dataset), and computed the Pearson correlation between the predicted and actual response in the test dataset. The distribution of correlation coefficients computed *with* shuffling (i.e., null distribution) was eventually compared to the one *without* shuffling (i.e., observed response). If the correlation coefficient of the observed response was greater than 95% of the correlation coefficients from the null distribution, this axis-coding model was considered *significant*. The correlation coefficient could also indicate the model’s predictability and thus be compared between different units.

### Data analysis: selection of attention-selective units

We used the mean firing rate in a time window of 150 ms to 225 ms after fixation onset as the response to each fixation. For each unit, if there was a significant difference in response (determined using a two-tailed Wilcoxon signed-rank test, with a significance threshold of P < 0.05) between fixations on targets and fixations on distractors, it was classified as an *attention-selective unit*. Similarly, for V4 units with a focal peripheral RF (as described above), we compared the response between targets and distractors within the RF in the same time window as for foveal units. Lastly, we calculated the attentional effect as the difference in firing rate between the same stimuli when they served as targets versus distractors.

### Data analysis: representational distance

We used the mean firing rate in a time window of 50 ms to 150 ms after fixation onset as the response to each fixation. For a population of units, we calculated the Euclidean distance between units. Change in population geometry was described using the cosine angle between the neuronal vectors: 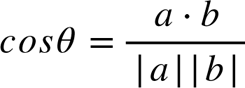, where *a* and *b* are the neuronal vectors for different conditions.

### Data analysis: spike-LFP coherence

We implemented the spike-LFP coherence analysis using the Chronux toolbox (www.chronux.org) in MATLAB. We used a single Hanning taper across frequencies, but we derived similar results using multitaper methods for higher frequencies (> 25 Hz) [56]. Coherence between two signals, *x* and *y*, was calculated using the following formula:

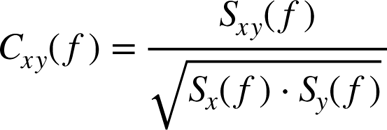

where *S_x_*(*f*) and *S_y_*(*f*) denote the auto-spectra and *S_xy_*(*f*) represents the cross-spectrum of the two signals *x* and *y*. Auto-spectra and cross-spectra were averaged across trials before coherence calculation. We used a 200-ms time window for each fixation (0 to 200 ms relative to fixation onset). Notably, we used an equal number of fixations and an equal number of spikes between conditions to calculate coherence for a given pair of recording sites, thus eliminating bias from different sample sizes. To avoid spikes contributing to the LFP recorded on the same electrode, we utilized signals from two different electrodes to calculate coherence. Furthermore, we did not select LFPs based on their category selectivity, attention selectivity, or the selectivity of the associated units (e.g., the LFP signals could come from contacts with both face-selective units and house-selective units).

### Data analysis: Granger causality

We utilized the open-source MATLAB toolbox “Granger causal connectivity analysis” (GCCA) [57] for our study. Frequency-domain Granger causalities were calculated during the same period as in coherence analysis between spikes and LFP across various brain areas. Two preprocessing steps, namely detrending and demeaning, were applied to the LFPs. The GCCA toolbox functions “cca_detrend” and “cca_rm_ensemblemean” were used to subtract the best-fitting line of the LFP for each fixation and the ensemble mean of the LFP, respectively. Subsequently, we employed a KPSS test to assess the stationarity of the LFP data after preprocessing, and any non-stationary LFP data were excluded from the analysis. Frequency-domain Granger causality was calculated based on the stationary LFPs after preprocessing using the function “cca_pwcausal” from the toolbox. The calculation is as follows:

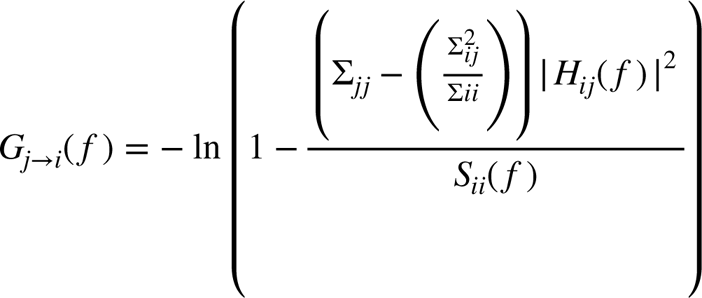

where *S_ii_*(*f*) is the power spectrum of variable *i* at frequency *f*, *H* is the transfer matrix, and *Σ* is the noise covariance matrix [57].

---

**Fig. S1.**
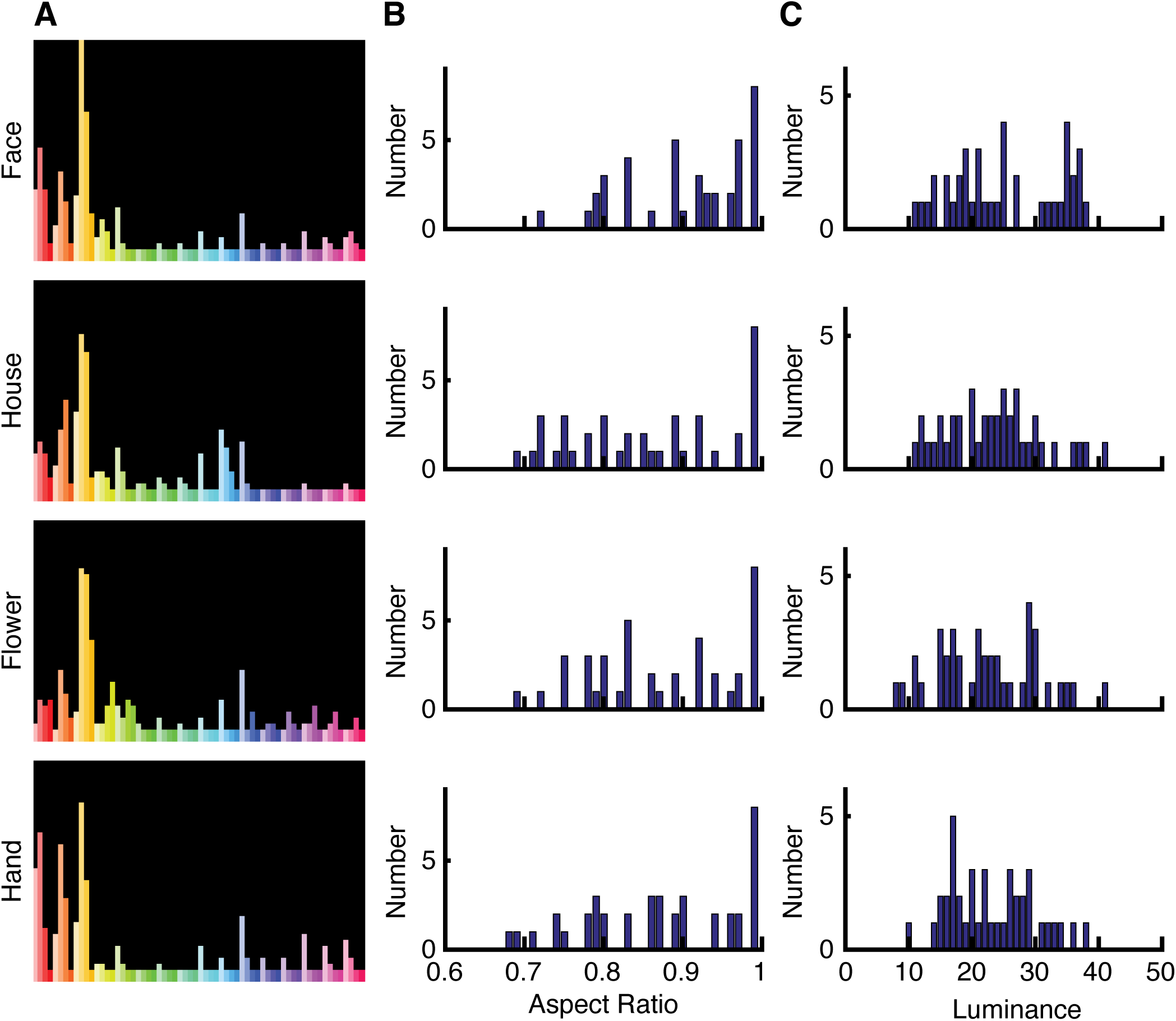
Characterization of the stimuli. **(A)** The stimuli from different categories did not exhibit significant differences in pixel-wise hue and saturation (χ^2^-test: P > 0.05). **(B)** The stimuli from different categories did not exhibit significant differences in aspect ratio (P > 0.05). **(C)** The stimuli from different categories did not exhibit significant differences in luminance (P > 0.05). Shown are histograms for each stimulus category.

**Fig. S2.**
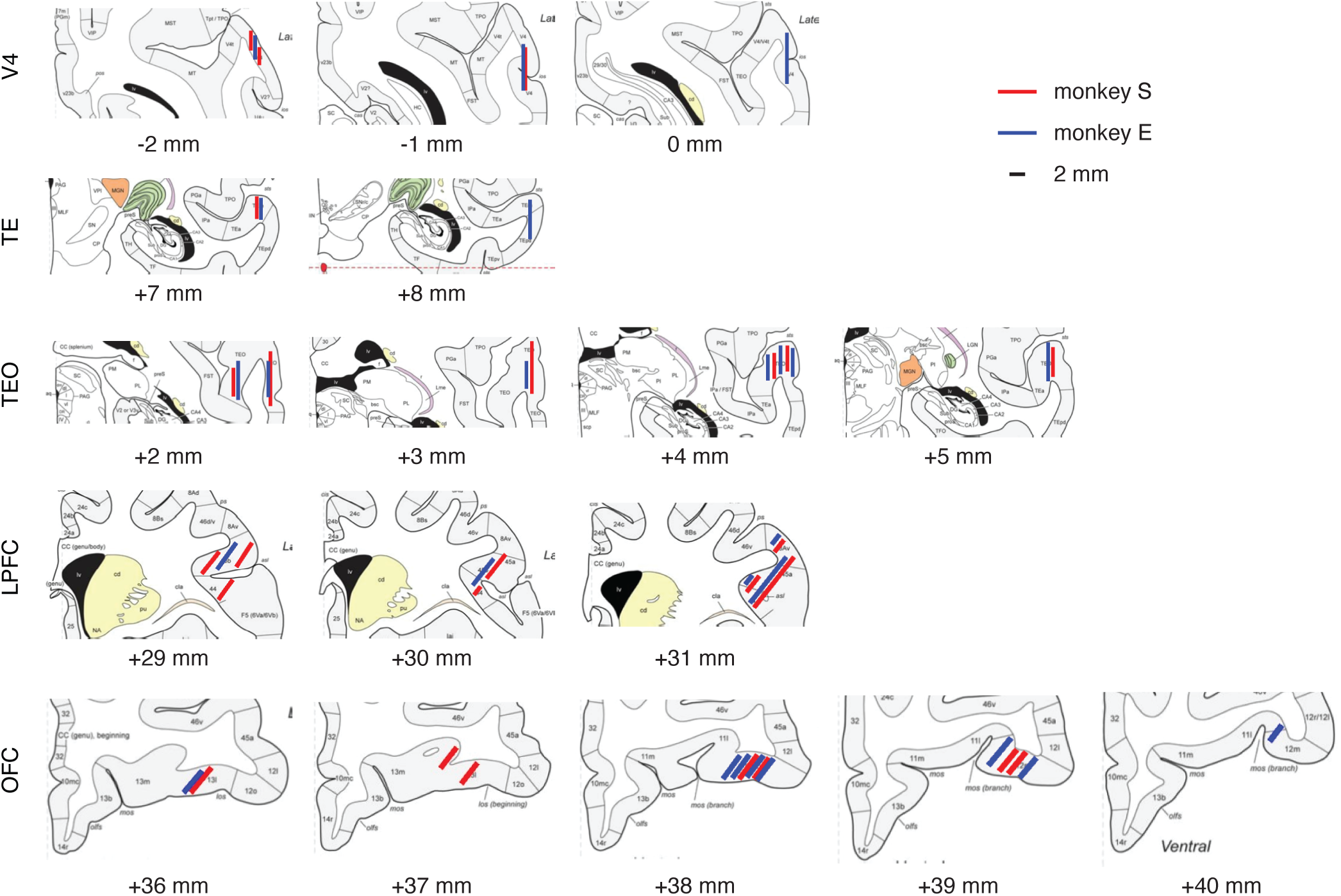
Recording sites in two monkeys overlaid on the atlas of the rhesus monkey brain in stereotaxic coordinates [58]. **(A)** V4. **(B)** TE. **(C)** TEO. **(D)** LPFC. **(E)** OFC. The red and blue lines represent the estimated spatial range of recordings in monkey S and monkey E, respectively. The numbers below indicate the rostral (+) or caudal (-) distances of the slices from the infraorbital ridge (Ear Bar Zero).

**Fig. S3.**
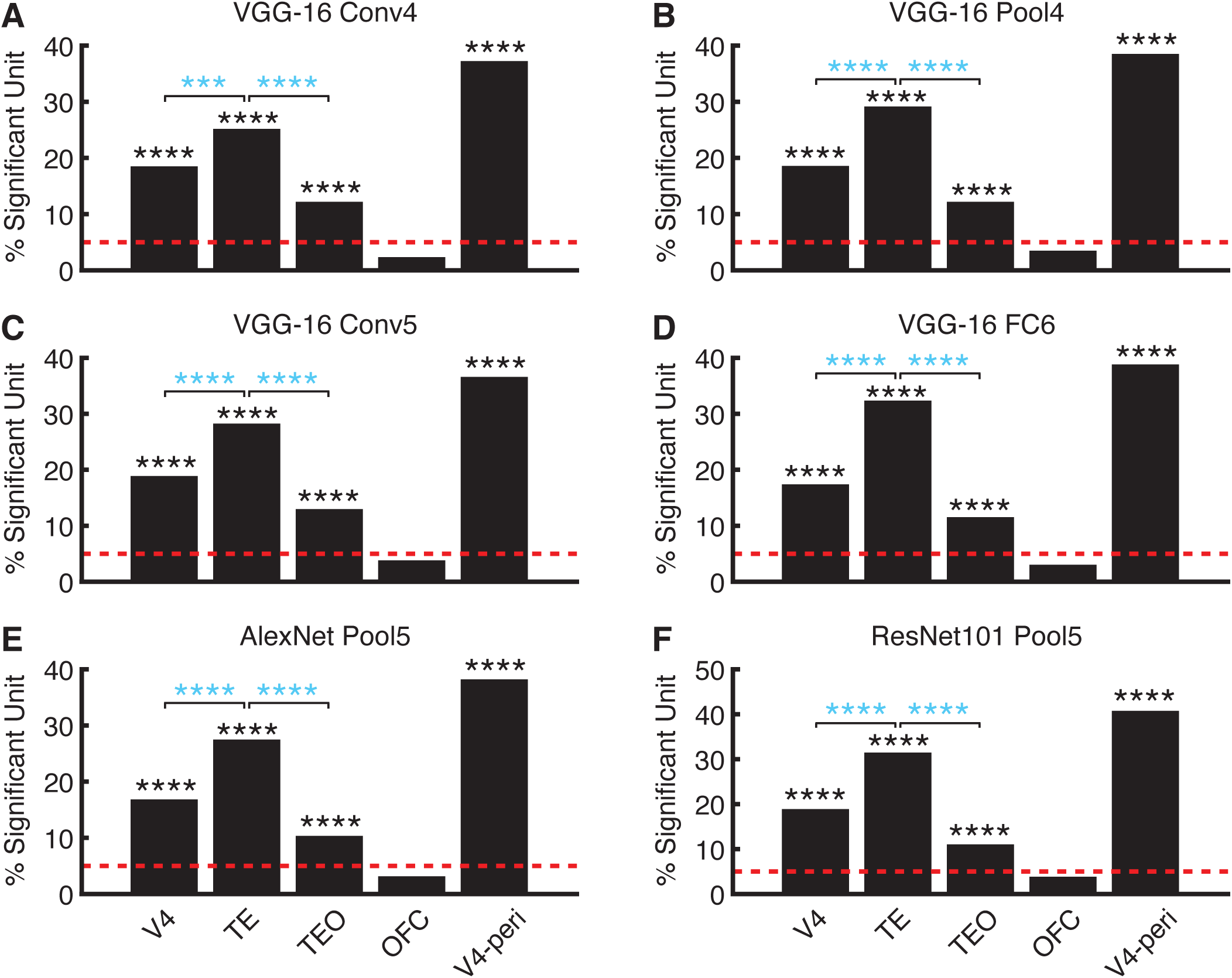
Control analysis for axis coding. **(A-D)** The proportion of axis-coding units for each brain area using features from different VGG-16 layers. **(E)** The proportion of axis-coding units for each brain area using features from the AlexNet. **(F)** The proportion of axis-coding units for each brain area using features from the ResNet. Black asterisks indicate a significant above-chance (5%) number of units (binomial test). Blue asterisks indicate a significant difference between brain areas (χ^2^-test). ****: P < 0.0001.

**Fig. S4.**
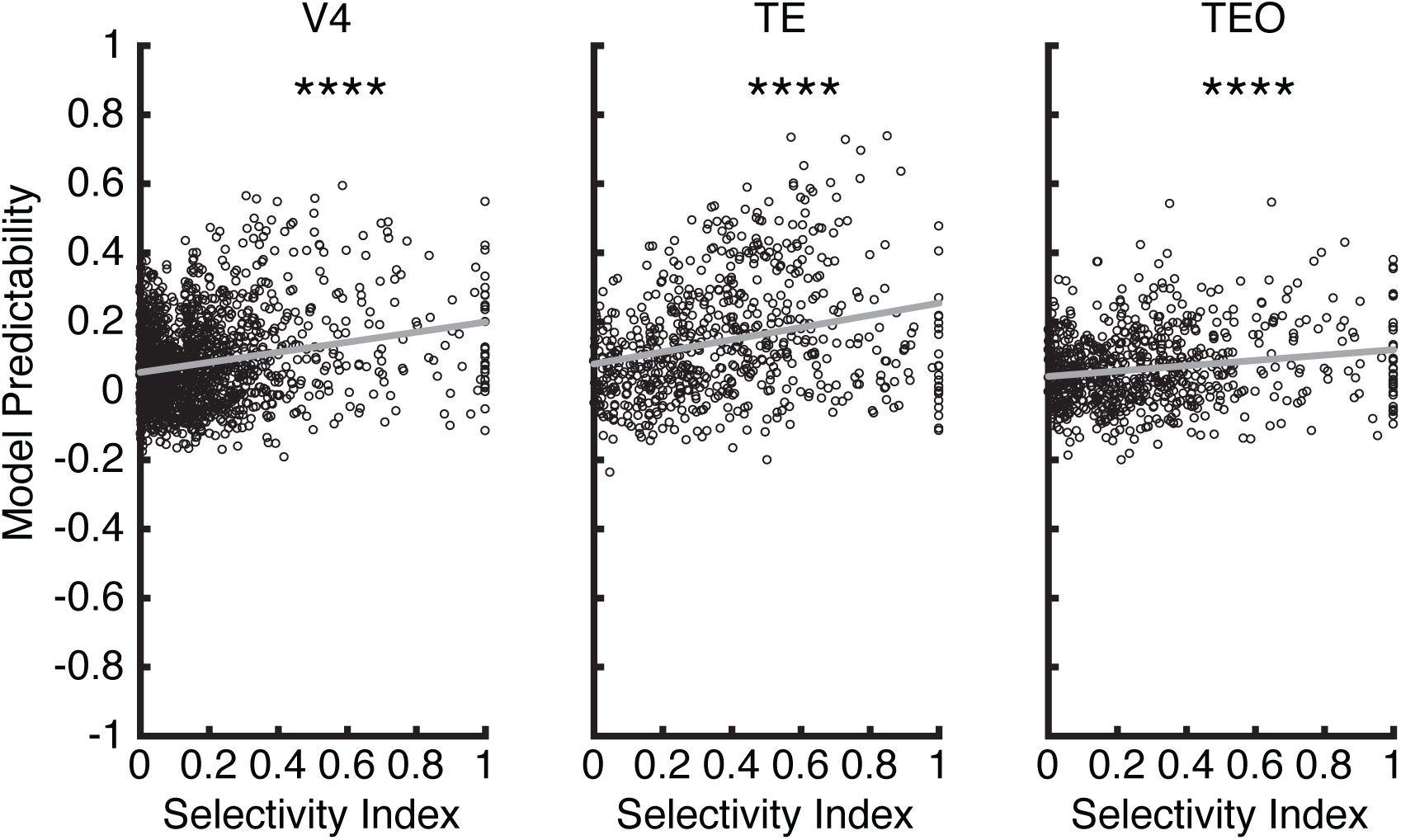
Correlation between axis coding and visual category selectivity. Axis coding is indexed by model predictability (see **Methods**). Each dot represents a unit. Asterisks indicate a significant Pearson correlation. ****: P < 0.0001.

**Fig. S5.**
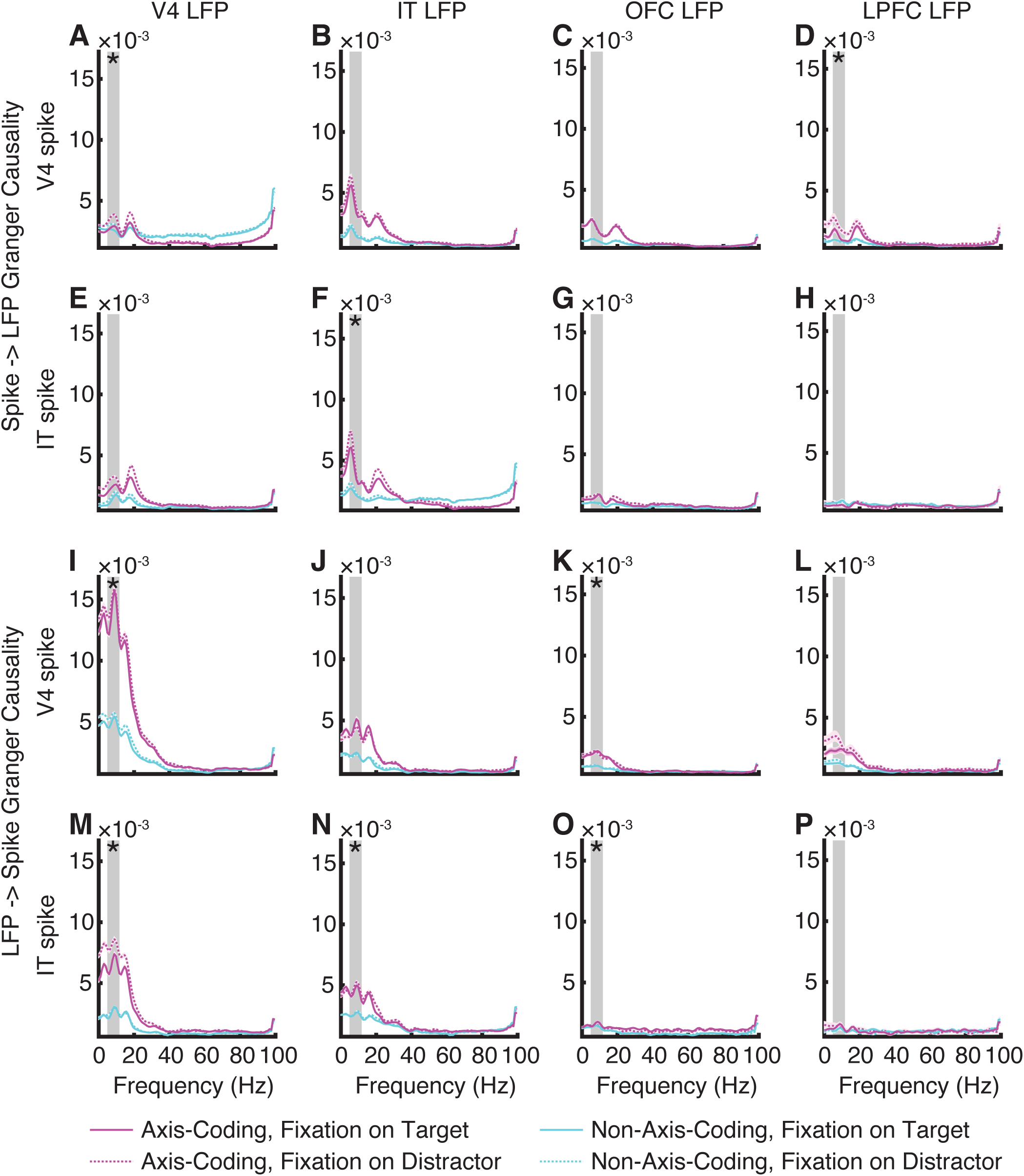
Granger causality. **(A)** V4 spike influence on V4 LFP. **(B)** V4 spike influence on IT LFP. **(C)** V4 spike influence on OFC LFP. **(D)** V4 spike influence on LPFC LFP. **(E)** IT spike influence on V4 LFP. **(F)** IT spike influence on IT LFP. **(G)** IT spike influence on OFC LFP. **(H)** IT spike influence on LPFC LFP. **(I)** V4 LFP influence on V4 spike. **(J)** IT LFP influence on V4 spike. **(K)** OFC LFP influence on V4 spike. **(L)** LPFC LFP influence on V4 spike. **(M)** V4 LFP influence on IT spike. **(N)** IT LFP influence on IT spike. **(O)** OFC LFP influence on IT spike. **(P)** LPFC LFP influence on IT spike. Legend conventions as in Fig. 6.

## References

1. Kastner, S. and L.G. Ungerleider, Mechanisms of visual attention in the human cortex, in Annual Review of Neuroscience. 2000. p. 315–341.

2. Corbetta, M. and G.L. Shulman, Control of goal-directed and stimulus-driven attention in the brain. Nat Rev Neurosci, 2002. 3(3): p. 201–15.

3. Petersen, S.E. and M.I. Posner, The Attention System of the Human Brain: 20 Years After. Annual Review of Neuroscience, 2012. 35(1): p. 73–89.

4. Moore, T. and M. Zirnsak, Neural Mechanisms of Selective Visual Attention. Annual Review of Psychology, 2017. 68(1): p. 47–72.

5. Fiebelkorn, I.C. and S. Kastner, Functional Specialization in the Attention Network. Annual Review of Psychology, 2020. 71(1): p. 221–249.

6. Chelazzi, L., et al., A neural basis for visual search in inferior temporal cortex. Nature, 1993. 363(6427): p. 345–347.

7. Buschman, T.J. and E.K. Miller, Top-Down Versus Bottom-Up Control of Attention in the Prefrontal and Posterior Parietal Cortices. Science, 2007. 315(5820): p. 1860–1862.

8. Zhou, H., Robert J. Schafer, and R. Desimone, Pulvinar-Cortex Interactions in Vision and Attention. Neuron, 2016. 89(1): p. 209–220.

9. Buschman, Timothy J. and S. Kastner, From Behavior to Neural Dynamics: An Integrated Theory of Attention. Neuron, 2015. 88(1): p. 127–144.

10. Logothetis, N.K. and D.L. Sheinberg, Visual object recognition. Annual Review of Neuroscience, 1996. 19(1): p. 577–621.

11. Gross, C.G., How Inferior Temporal Cortex Became a Visual Area. Cerebral Cortex, 1994. 4(5): p. 455–469.

12. Tanaka, K., Mechanisms of visual object recognition: monkey and human studies. Current Opinion in Neurobiology, 1997. 7(4): p. 523–529.

13. Freeman, W.J., Mass action in the nervous system. Vol. 2004. 1975: Citeseer.

14. Hinton, G.E., Distributed representations. 1984.

15. Rolls, E.T., A. Treves, and M.J. Tovee, The representational capacity of the distributed encoding of information provided by populations of neurons in primate temporal visual cortex. Experimental Brain Research, 1997. 114(1): p. 149–162.

16. Churchland, P.S. and T.J. Sejnowski, The computational brain. 2016: MIT press.

17. Chang, L. and D.Y. Tsao, The Code for Facial Identity in the Primate Brain. Cell, 2017. 169(6): p. 1013–1028.e14.

18. Bashivan, P., K. Kar, and J.J. DiCarlo, Neural population control via deep image synthesis. Science, 2019. 364(6439): p. eaav9436.

19. Ponce, C.R., et al., Evolving Images for Visual Neurons Using a Deep Generative Network Reveals Coding Principles and Neuronal Preferences. Cell, 2019. 177(4): p. 999–1009.e10.

20. Bao, P., et al., A map of object space in primate inferotemporal cortex. Nature, 2020. 583(7814): p. 103–108.

21. Rutishauser, U., et al., The Architecture of Human Memory: Insights from Human Single-Neuron Recordings. The Journal of Neuroscience, 2021. 41(5): p. 883.

22. Sheinberg, D.L. and N.K. Logothetis, Noticing Familiar Objects in Real World Scenes: The Role of Temporal Cortical Neurons in Natural Vision. The Journal of Neuroscience, 2001. 21(4): p. 1340–1350.

23. Bichot, N.P., A.F. Rossi, and R. Desimone, Parallel and Serial Neural Mechanisms for Visual Search in Macaque Area V4. Science, 2005. 308(5721): p. 529–534.

24. Zhou, H. and R. Desimone, Feature-Based Attention in the Frontal Eye Field and Area V4 during Visual Search. Neuron, 2011. 70(6): p. 1205–1217.

25. Chelazzi, L., Neural mechanisms for stimulus selection in cortical areas of the macaque subserving object vision. Behavioural Brain Research, 1995. 71(1–2): p. 125–134.

26. Mazer, J.A. and J.L. Gallant, Goal-Related Activity in V4 during Free Viewing Visual Search: Evidence for a Ventral Stream Visual Salience Map. Neuron, 2003. 40(6): p. 1241–1250.

27. Burrows, B.E. and T. Moore, Influence and Limitations of Popout in the Selection of Salient Visual Stimuli by Area V4 Neurons. The Journal of Neuroscience, 2009. 29(48): p. 15169–15177.

28. Zhang, J., et al., Visual Attention in The Fovea and The Periphery during Visual Search. bioRxiv, 2021: p. 2021.11.22.469359.

29. Yamins, D.L.K., et al., Performance-optimized hierarchical models predict neural responses in higher visual cortex. Proceedings of the National Academy of Sciences, 2014. 111(23): p. 8619.

30. Kriegeskorte, N., M. Mur, and P. Bandettini, Representational similarity analysis - connecting the branches of systems neuroscience. Frontiers in Systems Neuroscience, 2008. 2(4).

31. Cao, R., et al., Neural mechanisms of face familiarity and learning in the human amygdala and hippocampus. Cell Reports, 2024. 43(1): p. 113520.

32. Gothard, K.M., Multidimensional processing in the amygdala. Nature Reviews Neuroscience, 2020. 21(10): p. 565–575.

33. Wang, S., et al., Encoding of Target Detection during Visual Search by Single Neurons in the Human Brain. Current Biology, 2018. 28(13): p. 2058–2069.e4.

34. Maunsell, J.H.R. and S. Treue, Feature-based attention in visual cortex. Trends in Neurosciences, 2006. 29(6): p. 317–322.

35. Ruff, D.A. and M.R. Cohen, Simultaneous multi-area recordings suggest that attention improves performance by reshaping stimulus representations. Nature Neuroscience, 2019. 22(10): p. 1669–1676.

36. Yassa, M.A. and C.E.L. Stark, Pattern separation in the hippocampus. Trends in Neurosciences, 2011. 34(10): p. 515–525.

37. Leal, S.L. and M.A. Yassa, Integrating new findings and examining clinical applications of pattern separation. Nature Neuroscience, 2018. 21(2): p. 163–173.

38. Freedman, D.J., et al., Experience-Dependent Sharpening of Visual Shape Selectivity in Inferior Temporal Cortex. Cerebral Cortex, 2005. 16(11): p. 1631–1644.

39. Anderson, B., et al., Effects of Familiarity on Neural Activity in Monkey Inferior Temporal Lobe. Cerebral Cortex, 2008. 18(11): p. 2540–2552.

40. Fries, P., et al., Modulation of Oscillatory Neuronal Synchronization by Selective Visual Attention. Science, 2001. 291(5508): p. 1560–1563.

41. Yan, T. and H. Zhou, Synchronization between frontal eye field and area V4 during free-gaze visual search. Zoological Research, 2019. 40(5): p. 394.

42. Gong, X., W. Li, and H. Liang, Spike-field Granger causality for hybrid neural data analysis. Journal of Neurophysiology, 2019. 122(2): p. 809–822.

43. Wang, S., et al., Abstract goal representation in visual search by neurons in the human pre-supplementary motor area. Brain, 2019. 142(11): p. 3530–3549.

44. Baldauf, D. and R. Desimone, Neural Mechanisms of Object-Based Attention. Science, 2014. 344(6182): p. 424–427.

45. Buschman, Timothy J., et al., Synchronous Oscillatory Neural Ensembles for Rules in the Prefrontal Cortex. Neuron, 2012. 76(4): p. 838–846.

46. Fiebelkorn, I.C. and S. Kastner, A Rhythmic Theory of Attention. Trends in Cognitive Sciences, 2019. 23(2): p. 87–101.

47. Thompson, K.G., et al., Perceptual and motor processing stages identified in the activity of macaque frontal eye field neurons during visual search. Journal of Neurophysiology, 1996. 76(6): p. 4040–4055.

48. Freiwald, W.A., D.Y. Tsao, and M.S. Livingstone, A face feature space in the macaque temporal lobe. Nat Neurosci, 2009. 12(9): p. 1187–1196.

49. Freiwald, W.A. and D.Y. Tsao, Functional Compartmentalization and Viewpoint Generalization Within the Macaque Face-Processing System. Science, 2010. 330(6005): p. 845.

50. Simonyan, K. and A. Zisserman. Very deep convolutional networks for large-scale image recognition. in International Conference on Learning Representations. 2015.

51. Krizhevsky, A., I. Sutskever, and G.E. Hinton, ImageNet classification with deep convolutional neural networks, in Proceedings of the 25th International Conference on Neural Information Processing Systems - Volume 1. 2012, Curran Associates Inc.: Lake Tahoe, Nevada. p. 1097–1105.

52. He, K., et al. Deep residual learning for image recognition. in Proceedings of the IEEE conference on computer vision and pattern recognition. 2016.

53. Cao, R., et al., Feature-based encoding of face identity by single neurons in the human medial temporal lobe. bioRxiv, 2020: p. 2020.09.01.278283.

54. Kar, K., et al., Evidence that recurrent circuits are critical to the ventral stream’s execution of core object recognition behavior. Nature Neuroscience, 2019. 22(6): p. 974–983.

55. Kar, K. and J.J. DiCarlo, Fast Recurrent Processing via Ventrolateral Prefrontal Cortex Is Needed by the Primate Ventral Stream for Robust Core Visual Object Recognition. Neuron, 2021. 109(1): p. 164–176.e5.

56. Fries, P., et al., The Effects of Visual Stimulation and Selective Visual Attention on Rhythmic Neuronal Synchronization in Macaque Area V4. The Journal of Neuroscience, 2008. 28(18): p. 4823.

57. Seth, A.K., A MATLAB toolbox for Granger causal connectivity analysis. Journal of Neuroscience Methods, 2010. 186(2): p. 262–273.

58. Saleem, K.S. and N.K. Logothetis, A combined MRI and histology atlas of the rhesus monkey brain in stereotaxic coordinates. 2012, London: Academic Press.

